# Infant CD4 T-cell response to SARS-CoV-2 mRNA vaccination is restricted in cytokine production and modified by vaccine manufacturer

**DOI:** 10.1101/2025.04.02.646864

**Authors:** M Quinn Peters, Amber L. Young, Jennifer E. Stolarczuk, Madeline Glad, Erik Layton, Jennifer K. Logue, Nana K. Minkah, Helen Y. Chu, Janet A. Englund, D. Noah Sather, Chetan Seshadri, Alisa Kachikis, Whitney E. Harrington

## Abstract

**BACKGROUND:** Safe and effective vaccines are a key preventative measure to protect infants from SARS-CoV-2 infection and disease. Although mRNA vaccines induce robust antibody titers in infants, little is known about the quality of CD4 T-cell responses induced by vaccination. CD4 T-cell responses are important in orchestrating coordinated immune responses during infection and may help to limit disease severity. METHODS: To characterize the CD4 T-cell response to SARS-CoV-2 mRNA vaccination in infants, we sampled blood from 13 infants before and after primary SARS-CoV-2 mRNA vaccine series; samples from 12 historical vaccinated adults were used for comparisons. PBMC were stimulated with Spike peptide pools and the ability of CD4 T-cells to secrete Th1, Th2, and Th17 cytokines was quantified. A measure of polyfunctionality was generated using the COMPASS algorithm. RESULTS: We observed a significant increase in CD4 T-cells producing IL-2 (0.01% vs. 0.08%, p=0.04) and TNF-α (0.007% vs. 0.07%, p=0.007) following vaccination in infants but a more muted induction of IFN-γ production (0.01% vs 0.04%, p=0.08). This contrasted with adults, in whom vaccination induced robust production of IFN-γ, IL-2, and TNF-α. Th2 and Th17 responses were limited in both infants and adults. In infants, CD4 T-cell responses post-vaccination were greater in those who received mRNA-1273 versus BNT162b. In contrast to CD4 T-cell responses, Spike-specific IgG titers were similar in infants and adults. CONCLUSIONS: These data suggest that infants have restricted induction of cytokine producing CD4 T-cells following SARS-CoV-2 mRNA vaccination relative to adults.

## INTRODUCTION

Vaccination against SARS-CoV-2 is a critical intervention for prevention of morbidity and mortality associated with COVID-19 disease. The widely available mRNA-based vaccines BNT162b (Pfizer-BioNTech) and mRNA-1273 (Moderna) were initially developed for adults but were subsequently adapted for pediatric populations with altered antigen amount and vaccine schedule and were approved for infants over 6 months of age by the US Food and Drug Administration in June 2022 (1). Immunogenicity comparison studies indicate that the mRNA vaccines induced comparable anti-Spike IgG titers in infants and adults (2, 3), but efficacy against symptomatic COVID-19 was only 75.8% in children 6 months to 2 years for BNT162b and 50.6% for mRNA-1273 (2, 3), lower than early efficacy studies in adults (4, 5).

Generation of Spike-specific antibodies, and in particular those that neutralize virus, have been the predominant focus of vaccine efforts (3, 6, 7), as antibodies are critical to preventing infection (8). However, once infected, effective viral clearance and control of disease severity may be primarily T-cell, particularly Th1, mediated (9-14). T-cell responses may also be less susceptible to immune escape, relative to antibodies (13, 15). In adults, prior studies have identified both CD4 and CD8 T-cell responses following SARS-CoV-2 mRNA vaccination, though CD4 responses are predominant (15-18). In contrast, few prior studies have investigated T-cell responses generated by SARS2 mRNA vaccines in infants (19). This may reflect the challenge in obtaining sufficient whole blood from infants to conduct T-cell stimulation assays, a problem compounded by the low frequency of antigen-specific T-cells generated by vaccination (20, 21).

To address this critical gap in knowledge, we characterized the infant CD4 T-cell response to primary SARS-CoV-2 mRNA vaccination and used a group of adult vaccine recipients as reference. We interrogated the frequency of Spike-specific CD4 T-cells as well as their functionality.

## MATERIAL AND METHODS

### Cohort

Maternal-infant dyads were enrolled into the “Maternal immunizations in low- and high-risk pregnancies” (MATIMM) study approved by the University of Washington Institutional Review Board (STUDY00008491); for infant participation, informed consent was obtained from mothers. Families who intended to vaccinate their infant <12 months of age with a SARS-CoV-2 mRNA vaccine (BNT162b or mRNA-1273) were identified. Peripheral blood was collected from infants prior to the first dose of the vaccine and again 3-7 weeks following completion of the primary series (3^rd^ dose for BNT162b and 2^nd^ dose for mRNA-1273). Whole blood from EDTA vacutainers was processed for plasma and PBMC as previously described (22). Clinical data were abstracted from patient charts, including dates of vaccination and vaccine manufacturer.

For adult controls, plasma and PBMC from adults vaccinated with either the BNT162b or mRNA-1273 primary series previously banked by the HAARVI Study (UW: STUDY00000959) (23), SCRI SARS2 Vaccine Study (SC: STUDY00003064), or the SCRI CGIDR Biorepository (SC: STUDY00002048) were utilized (24). For all adult control samples, informed consent was obtained into the parent study and subsequent use of the banked de-identified samples was included within the IRB-approved protocol. Samples were collected before vaccination and then 2-7 weeks following completion of primary series. Adults were selected to include an equal distribution of BNT162b and mRNA-1273 vaccinated individuals.

Plasma for all adult samples and all but one infant pre-vaccine sample was tested for SARS-CoV-2 nucleocapsid (NC) IgG by enzyme linked immunosorbent assay (detailed below). For the single exception, plasma at this timepoint for the infant had previously tested negative for NC-IgG and positive for Spike-IgG using Roche Diagnostics GmbH. Elecsys® Anti-SARS-CoV-2 2023.

### Intracellular Cytokine Staining

PBMCs were thawed at 37°C and transferred to prewarmed RPMI 1640 with l-glutamine (Thermo Fisher) with 10% FBS (Biowest) and 0.2% Benzonase (Millipore Sigma). Cells were then rested overnight at 37°C with 5% CO2. The next day, cells were enumerated, resuspended in media, and seeded into a 96-well U-bottom plate, split across three wells at a density of 1 to 2 million cells/well. S1 and S2 peptide pools optimized for CD4 stimulation containing 15mers with 11aa overlap (JPT Peptide Technologies) were prepared as previously described (25). Stimulation cocktails were prepared for the conditions of negative control: 0.2% DMSO (Sigma-Aldrich), Spike peptide pool (1 μg/ml of each S1 and S2), or positive control: 0.05 μg/ml phorbol-12-myristate-13-acetate and 0.7471 ug/ml Ionomycin. Stimulation cocktails also contained 10 μg/ml brefeldin A (Sigma-Aldrich), GolgiStop (BD Biosciences), 1 μg/ml CD28/49d (BD Biosciences, San Jose, CA), and anti-CD107a PE-Cy7 (BD Biosciences). Cells were stimulated for 6 hours at 37°C, followed by the addition of EDTA, and overnight storage at 4°C (20). Cells were subsequently stained sequentially with Fixable Aqua viability dye (Invitrogen) and anti-CCR7 (BD Biosciences), followed by RBC lysis with FACS Lyse (BD Biosciences). Cells were permeabilized FACS Perm II (BD Biosciences) and stained with an intracellular antibody cocktail (Supplemental Table 1), fixed with 1% paraformaldehyde (Electron Microscopy Solution), and then resuspended in PBS with 2 mM EDTA. Cells were stored at 4°C until acquisition.

Data was acquired on a BD LSR Fortessa (BD Biosciences) flow cytometer. Adult positive and negative biological controls were run in each experiment. The same control donors were used in each assay, and the pre and post vaccine timepoint samples from each individual were always run in the same experiment.

### Flow Cytometry Data Analysis

Flow cytometry data were analyzed using FlowJo v10.10 software. To compare monofunctional cytokine production, the background frequency of cells producing a cytokine in the DMSO condition was subtracted from that of the Spike peptide pool condition. Combinatorial Polyfunctionality Analysis of Ag-Specific T-cell Subsets (COMPASS; v1.32.0) (26) was used to analyze the functional profiles of antigen-specific T-cells in an unbiased and comprehensive manner. COMPASS outputs a functionality score (**FS**), which estimates the proportion of antigen-specific cell subsets, weighted by the degree of functionality. The FS was used to characterize the entire functional profile of a sample into one variable. COMPASS also outputs a polyfunctionality score (**PFS**), which summarizes the breath of polyfunctional responses to stimulation. Default settings were used to filter categories and samples with fewer than 3,000 CD4 T-cells were excluded from analysis. The R package ComplexHeatmap (v2.10.0) was used to visualize probabilities of response from the COMPASS output (27). One pre-vaccine infant sample did not meet the cell count threshold and was dropped from the analysis.

### Enzyme Linked Immunosorbent Assay

Plasma IgG titers to SARS-CoV-2 trimer and NC were determined using direct immobilization ELISA. Plasma was heat-inactivated for 1 hour at 56°C prior to the assay and centrifuged at 17,000 x g for 10 min. Immulon 2HB 96-well plates (Thermo Scientific, 3455) were coated with fifty nanograms per well of SARS-CoV-2 trimer or NC in 0.1M aHCO3, pH 9.5 overnight at room temperature. Plates were washed between each ELISA step with PBS containing 0.2% Tween-20. Coated plates were blocked with PBS, 10% non-fat milk, and 0.3% Tween-20. Following blocking, plasma samples were serially diluted over a range of 1:50 to 1:36,450 in PBS, 10% non-fat milk, 0.03% Tween-20 and incubated for 1 hour at 37°C. Bound antibodies were detected using goat anti-human IgG Fc-HRP (Southern Biotech, 2049-05) at 1:4000 dilution in PBS, 10% non-fat milk, 0.03% Tween-20. Plates were developed using 50 ul of TMB Peroxidase Substrate (SeraCare Life Sciences Inc, 5120-0083), then stopped after 3 minutes with 50 ul of 1N H2SO4. Absorbance at 450 nm was determined with the BioTek ELx800 microplate reader. Endpoint titers were defined as the reciprocal of plasma dilution at O.D. 0.1 after the subtraction of plate background. A negative serologic response was defined as < 1:50 for IgG.

### Statistical Analysis

Magnitude of background subtracted CD4 T-cell cytokine responses was compared between pre- and post-vaccine samples using paired T-tests. For comparison of post-vaccine response (individual cytokines, FS, PFS, and Spike-specific IgG) between infants and adults a multivariate regression model was utilized with adjustment for sex, time post-vaccination of blood sampling, and vaccine manufacturer. Similarly, for comparison of post-vaccine response (individual cytokines, FS, PFS, and Spike-specific IgG) by vaccine manufacturer a multivariate model was utilized separately for infants or adults with adjustment for sex, time post-vaccination of blood sampling, and age at vaccine series initiation. The infant who received a mixed vaccine series was excluded from this analysis. For comparison of endpoint titers, a value of 1:49 was used for samples negative at the initial 1:50 dilution. CD4 T-cell FS versus Spike endpoint titer was compared with Pearson’s correlation.

## RESULTS

### Cohort Characteristics

We enrolled 13 infants who received a primary series of either SARS-CoV-2 mRNA vaccine, which included one set of twins. There were four female and nine male infants. The median age of infants at the pre-vaccine timepoint was 6.9 months (range 6 to 11.5 months) (**Table 1**), which corresponded to a median of 1 day (range 0-16 days) prior to receipt of first vaccine dose. Six infants received the two-dose mRNA-1273 vaccine, six received the three-dose BNT162b vaccine, and one infant received two doses of BNT162b followed by one dose of mRNA-1273. Infants were a median of 10.1 (range: 7.9-15.1) months of age at the post-vaccine timepoint, which corresponded to a median of 28 days (range 22 to 42 days) post completion of primary vaccine series (**Figure 1**).

**Table 1.** Participant characteristics.

|  | Infants (n=13) | Adults (n=12) |
| --- | --- | --- |
| Infant sex, female, *n (%)* | 4 (31%) | 7 (58%) |
| Age at vaccine series initiation, *median (range)* | 6.9 (6.0-11.5) months | 37.5 (27-81) years |
| vaccine manufacturer, Pfizer, *n (%)* | 6 (46%) | 6 (50%) |
| Time between first blood draw and first vaccine dose (days), *median (range)* | 1 (0-16) days | 52 (2-323) days |
| Time between final vaccine dose and post-vaccine blood draw (days), *median (range)* | 28 (22-42) days | 14.5 (9-42) days |

**Figure 1.**
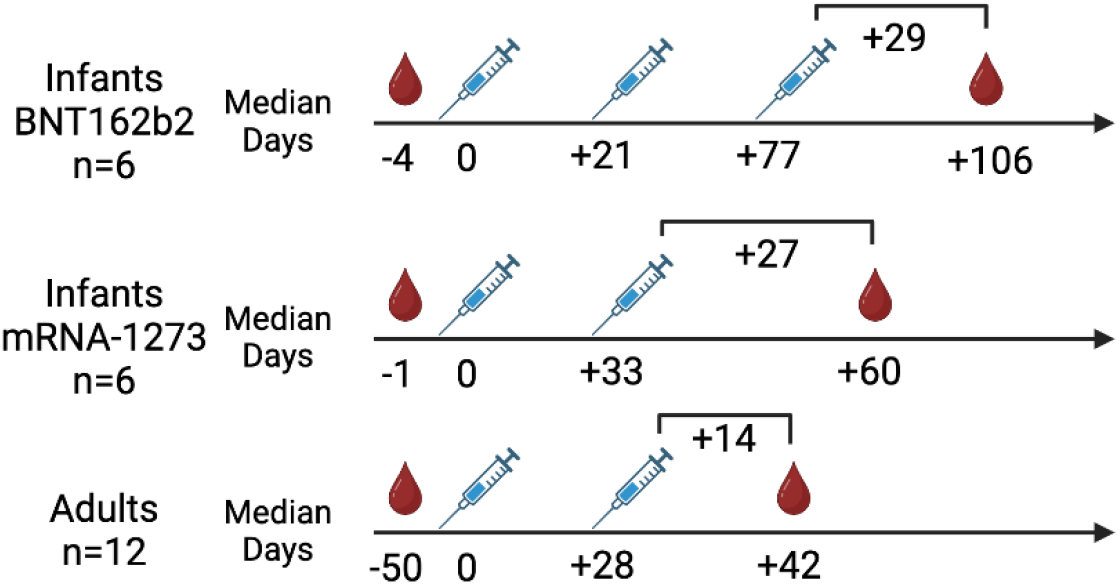
Vaccination overview. Infants (n=13) and adults (n=12) donated blood prior to SARS-Co-2 vaccination and again following vaccination. BNT162b2 primary series for infants included 3 doses; mRNA-1273 primary series for infants included 2 doses. One infant received two doses of BNT162b vaccine and one dose of the mRNA-1273 vaccine (not represented). Both adult primary series included 2 doses. Time points are relative to the first vaccine dose and represent medians. Created with BioRender.com.

The median age of adult controls was 37.5 years (range 27-81 years) (**Table 1**); there were seven female and five males. Six individuals received BNT162b and six received mRNA-1273. The pre-vaccine blood draw occurred a median of 52 days (range: 2-323 days) prior to the first dose of vaccine (**Figure 1**). The median time of blood drawn following the completion of primary series was 14.5 days (range 9 to 42 days).

### CD4 T-cell cytokine responses to SARS-COV-2 Spike following mRNA vaccination

Our primary aim was to characterize the infant CD4 T-cell response to the SARS-COV-2 mRNA vaccines. In the 13 infants sampled, there were universally low CD4 T-cell responses to Spike stimulation at the pre-vaccine timepoint. Between the pre- and post-vaccine timepoints, there were significant increases in the frequency of CD4 T-cells producing IL-2 (0.01% vs. 0.08%, p=0.04) and TNF-a (0.007% vs. 0.07%, p=0.007) in response to Spike peptide pool, but a more muted induction of IFNγ production (0.01% vs 0.04%, p=0.08). In addition, there were low frequencies of CD4 T-cells that produced Th2 cytokines IL-4, IL-5, and IL-13 (0.005% vs. 0.03%, p=0.07) or IL-17a (0.008% vs. 0.02%, p=0.2) in response to Spike stimulation, but these responses were not significantly different between pre- and post-vaccine timepoints (**Figure 2**). CD4 T-cells also significantly increased their expression of CD154 following stimulation with Spike peptide pool between pre- and post-vaccine timepoints (0.04% vs. 0.15%, p=0.04), but there was no change in expression of CD107a (0.03% vs. 0.05%, p=0.4).

**Figure 2.**
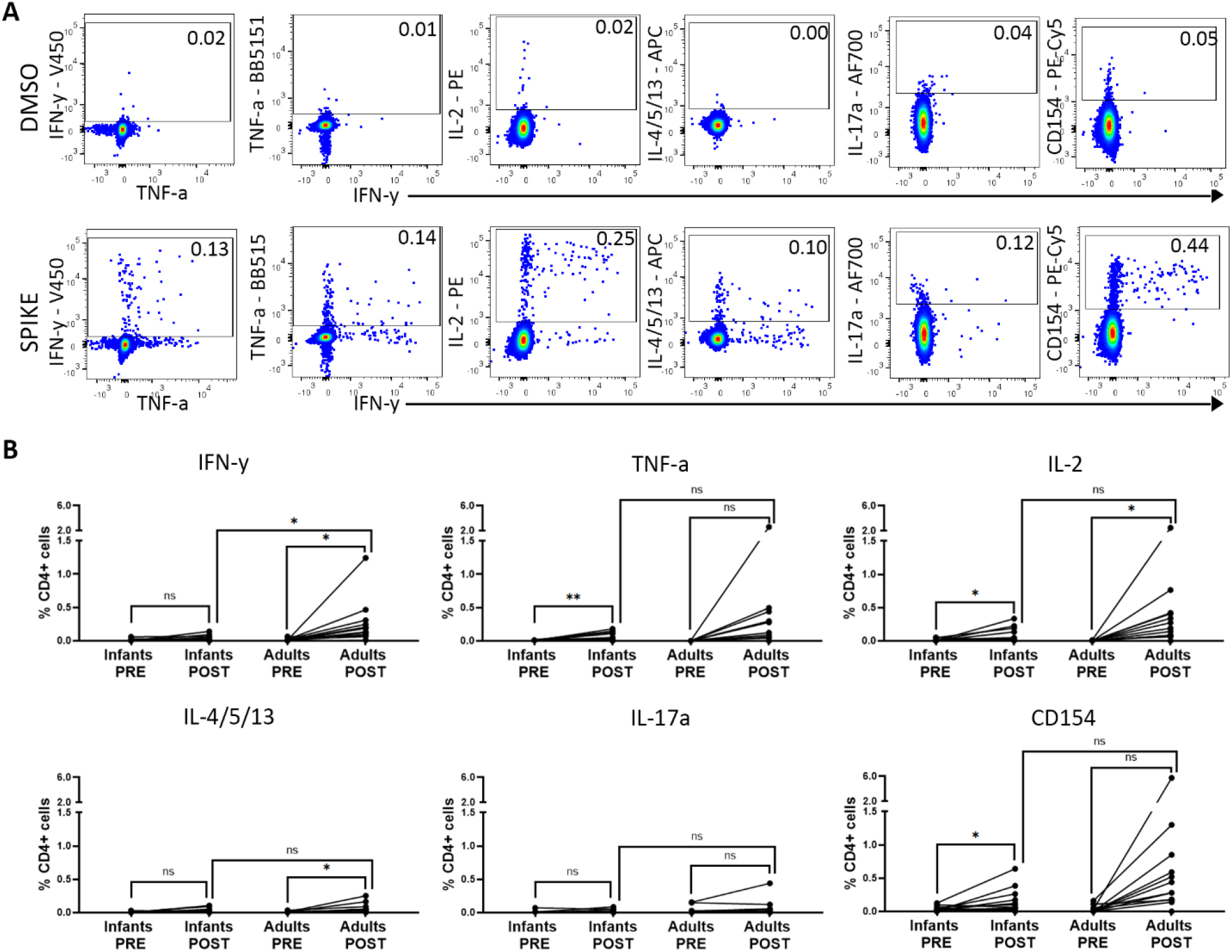
Cytokine production from CD4 T-cells before and after SARS2 mRNA vaccination. (A) Representative flow plots of infant CD4 T-cells expressing Th1, Th2, Th17 cytokines or CD154 in response to stimulation with DMSO or Spike peptide pool. (B) Summary plots of background subtracted frequencies of infant CD4 T-cells expressing cytokines in response to stimulation with Spike peptide pool. Background subtracted pre vs. post cytokine response compared with paired t tests. Post-vaccine response between adults and infants compared with multivariate regression model with adjustment for sex, time post-vaccination of blood sampling, and vaccine manufacturer. \**p* ≤ 0.05, \*\**p* ≤ 0.01.

In adults we observed a robust increase in CD4 T-cell production of all Th1 cytokines in response to Spike peptide pool between the pre- and post-vaccine timepoints (IFNγ: 0.02% vs. 0.26%, p=0.03; IL-2: 0.004% vs. 0.44%, p=0.05; TNF-a: 0.002% vs. 0.40%, p=0.08), though TNF-a was not statistically significant, consistent with prior reports (20). In addition, there was an increase in the frequency of CD4 T-cells producing Th2 cytokines (0.006% vs 0.06%, p=0.02), but not IL-17a (0% vs. 0.02%, p=0.2). There was a non-significant increase in the frequency of CD4 T-cells expressing CD154 (0.03% vs. 0.87%, p=0.09), and no change in the frequency of CD107a expressing cells (0.02% vs 0.03%, p=0.5).

At the post vaccine timepoint, IFNγ production was significantly higher among adults versus infants (adjusted difference: 0.26%, p=0.04). Production of TNF-α (adjusted difference: 0.40%, p=0.1), IL-2 (adjusted difference: 0.40%, p=0.1), and expression of CD154 (adjusted difference: 0.86%, p=0.1) were also higher, but did not reach significance (**Figure 2B**).

### CD4 T-cell polyfunctionality following SARS2 mRNA vaccination varies between infants and adults

To assess the functional diversity of vaccine-induced T-cell responses among infants comprehensively, we next used COMPASS to determine the probability of detecting a response greater than the background among all possible combinations of cytokine subsets. A response above background was detected in 20 out of 107 possible CD4 functional subsets, 15 of which were polyfunctional (**Figure 3A, B**). Several Spike-specific CD4 subsets were identified at pre-vaccine timepoints, which may be due to the cross-reactivity between SARS-CoV-2 Spike protein and other common human coronaviruses (28). Post vaccination, a polyfunctional subset producing canonical Th1 cytokines IFN-γ, IL-2, and TNF-α, in addition to CD154, was identified in 10 of 13 infant samples with moderate probability scores and 11 of 12 adult samples with high probability scores. Similarly, a polyfunctional subset positive for only IL-2, TNF-α, and CD154 was identified in all infant samples and 11 of 12 adult samples, and a subset with IFN-γ, IL-2, and CD154 was present in five infants, most with low probability scores, and all adult samples with high probability scores (**Figure 3B**). IL-17a was present in only two subsets: a monofunctional subset and a polyfunctional subset in combination with Th2 cytokines.

**Figure 3.**
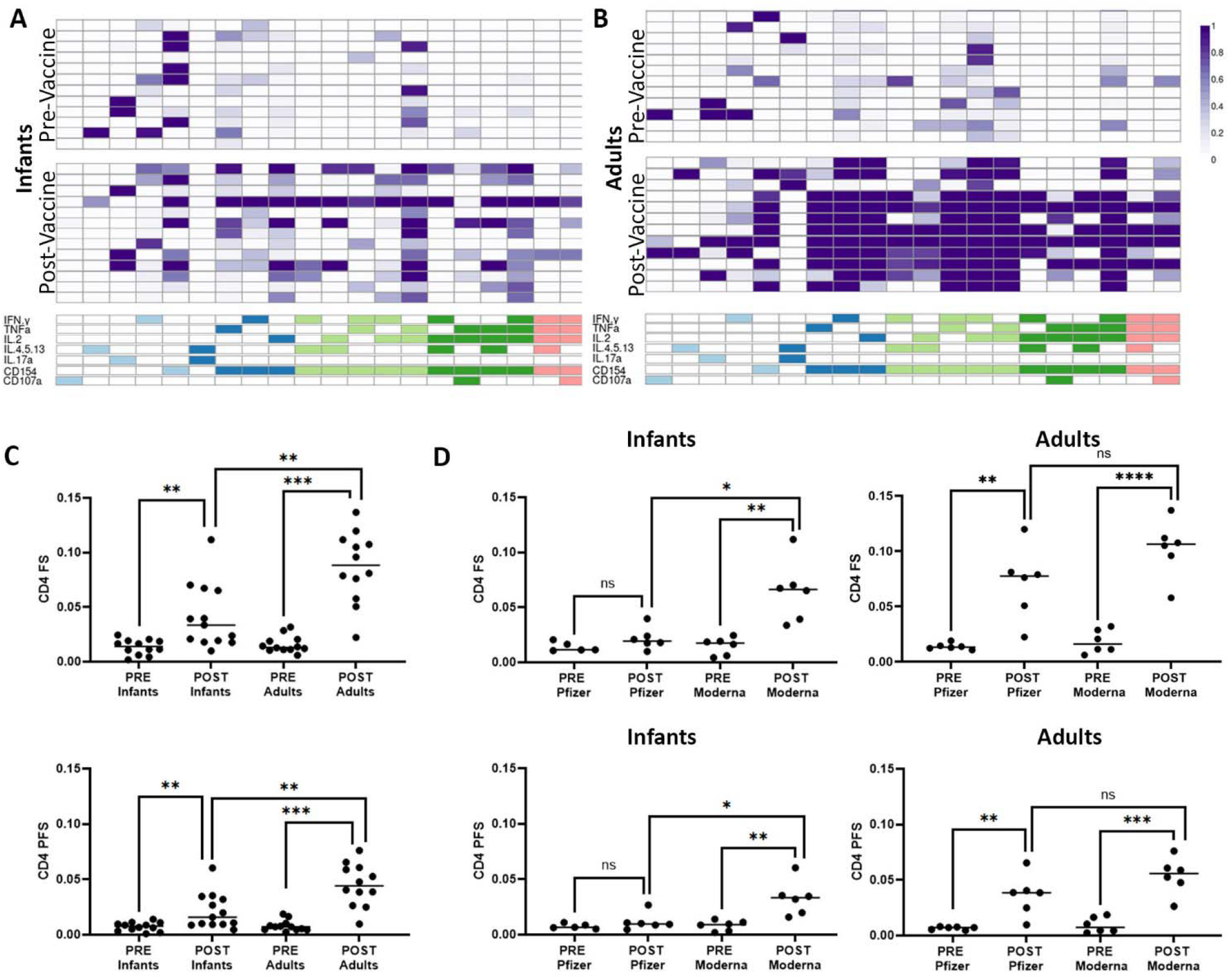
Functional breath of CD4 T-cell response determined by COMPASS analysis. (A) Results from COMPASS analysis of ICS data from infants displayed as a heatmap of probabilities. Rows represent individuals, with order matched between pre- and post-vaccine groups. Columns represent functional CD4 T-cell subsets identified in an unbiased manner by COMPASS. Columns of the same color share the same degree of functionality. (B) Probability heatmap output from COMPASS analysis of ICS data from adults. (C) FS and PFS by age group and vaccine status. (D) FS and PFS arranged by vaccine status and vaccine manufacturer for infants and adults. Post-vaccine response by vaccine manufacturer compared with multivariate regression with adjustment for sex, time post-vaccination of blood sampling, and age at vaccine series initiation, separately for adults or infants. \**p* < 0.05, \*\**p* < 0.01,***p<0.001.

Together, these data underscore the Th1 skew of CD4 T-cells in response to SARS-CoV-2 vaccination in both infants and adults, and the robust production of IFNγ in adults.

Both infants and adults showed significant increases in FS (infants: 0.01 to 0.04, p=0.009; adults: 0.02 to 0.09, p<0.001) and PFS (infants: 0.008 to 0.02, p=0.01; adults: 0.008 to 0.04, p<0.001) following vaccination. The FS and PFS of adults post-vaccination, however, was significantly higher than those of infants (FS adjusted delta: 0.04, p=0.005; PFS adjusted delta: 0.02, p=0.01) (**Figure 3C**).

### Accentuated induction of CD4 T-cell Spike-specific responses following mRNA-1273 versus BNT162b mRNA vaccination in infants

When considered by manufacturer, we observed a significantly higher frequency of post-vaccine CD4 T-cells producing IL-2 (adjusted delta: 0.14%, p=0.02) or Th2 cytokines (adjusted delta: 0.05%, p=0.02) and expressing CD154 (adjusted difference 0.22%, p=0.03) among infants vaccinated with mRNA-1273 versus BNT162b. There was also a trend toward increased IL-17 production (adjusted difference: 0.03% p=0.07) but no difference in TNF-α or INF-γ production or CD107a expression. Infants who received mRNA-1273 vs. BNT162b also had higher FS (adjusted difference: 0.05, p=0.02) and PFS (adjusted difference: 0.03, p=0.04) post-vaccination, driven by a lack of response in the BTN162b recipients (**Fig 3D)**. In adults, mRNA-1273 vs. BNT162b recipients had non-significantly higher frequency of CD4 T-cells producing IFN-γ (adjusted difference: 0.35%, p=0.1), TNF-α (adjusted difference: 0.69%, p=0.2), IL-2 (adjusted difference: 0.69%, p=0.2), and Th2 cytokines (adjusted difference: 0.08%, p=0.1) and significantly higher expression of CD154 (adjusted difference: 1.44%, p=0.02). There was no difference in production of IL-17 or expression of CD107a. Post-vaccine FS (adjusted difference: 0.03, p=0.1) and PFS (adjusted difference: 0.02, p=0.2) were also non-significantly increased in mRNA-1273 vs. BNT162b recipients (**Figure 3D**).

### Infants and adults generate similar Spike-specific IgG titers following vaccination

Prior work has reported similar magnitude of anti-Spike IgG induced following mRNA vaccination in adults and children aged 6 months to 5 years (2, 3). However, because of our observed differences in CD4 T-cell responses between adults and infants, we sought to confirm whether this was also true in our cohort. At the pre-vaccine timepoint, all adults were Spike-specific IgG negative, whereas in infants, five of 13 had detectable Spike-specific IgG titers (**Fig 4A**). Given that all infants were NC-specific IgG negative at the pre-vaccine timepoint, these titers likely reflect residual transplacental maternal IgG and not early infection. All infants and adults had robust induction of Spike-specific IgG (**Fig 4A, B**). At the post-vaccine timepoint, the mean endpoint titers amongst infants and adults were similar (mean log10 4.67 vs. 4.66; adjusted difference: log10 0.23, p=0.2). There was no difference in post-vaccine titer in infants who were positive vs. negative for Spike-specific IgG at the pre-vaccine timepoint (mean: log10 4.53 vs. 4.75; p=0.2). In children, there was no difference in post-vaccine Spike-specific IgG titer by vaccine manufacturer (adjusted difference: log10 −0.18, p=0.3); whereas, amongst adults, the titers were significantly higher in mRNA-1273 vs. BNT162b recipients (adjusted difference: log10 0.66, p=0.002). At the post-vaccine timepoint, there was no correlation between the magnitude of the Spike-specific T-cell FS and IgG endpoint titers (R^2^=0.08, p=0.8) (**Fig 4C**) in infants, whereas in adults there was a strong correlation (R^2^=0.81, p=0.001) (**Fig 4D**).

**Figure 4.**
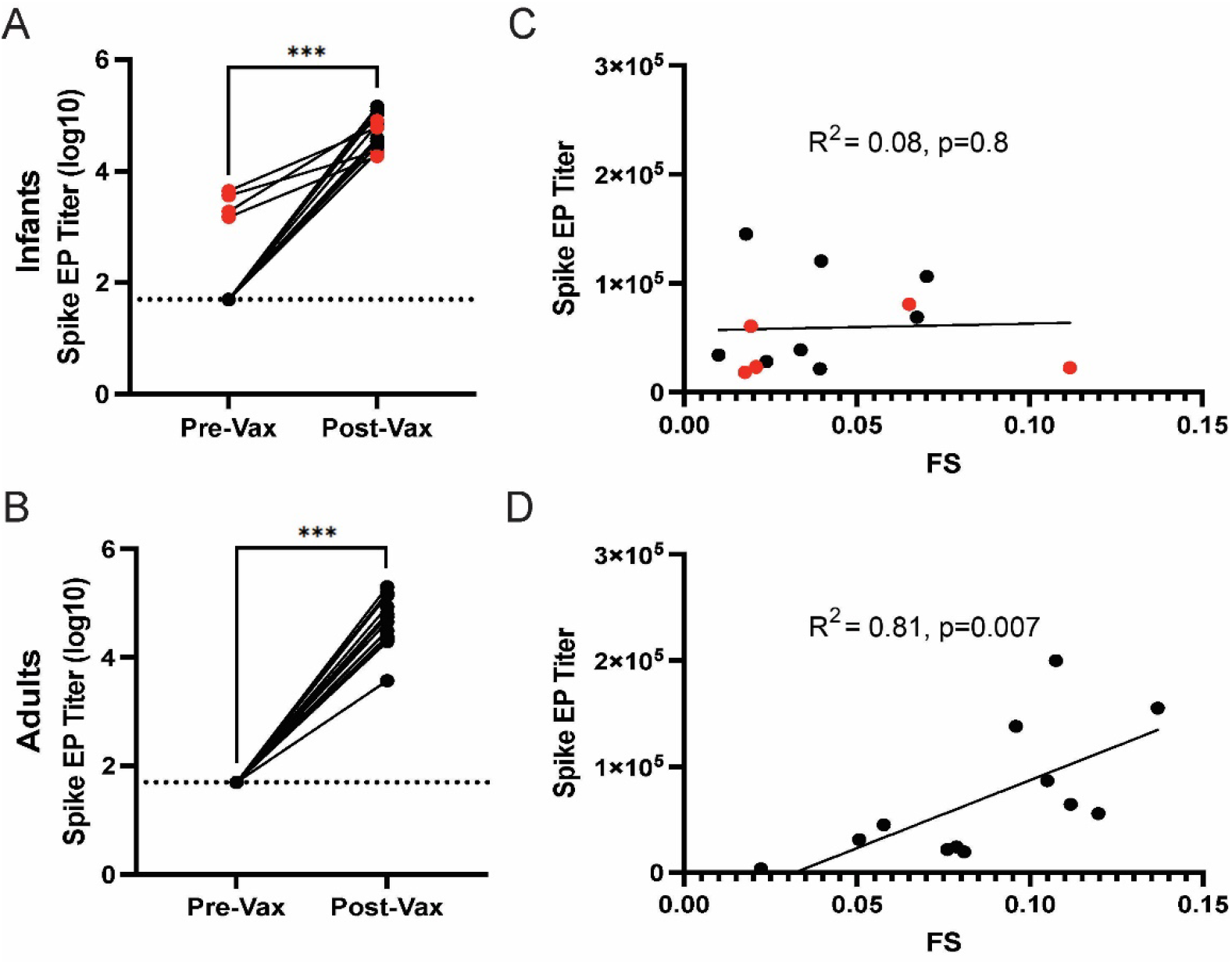
Anti-spike IgG antibody endpoint titers are a correlate of CD4 T-cell response following vaccination for adults but not infants. (A) Paired anti-spike IgG endpoint titers from infants and (B) adults before and after SARS-CoV-2 vaccination. All post-vaccine endpoint titers were above the positive threshold of log10 (50), represented as a dashed line. Infants with positive anti-Spike endpoint titers prior to vaccination (red) are likely due to transplacental maternal IgG transfer. (C) Infant and (D) adult anti-Spike IgG endpoint titers plotted against spike CD4 T-cell FS from post-vaccine time points, shown with a best-fit line. Pre-vaccine data ELISA not generated for one infant, but the sample was classified as positive for Spike IgG at the pre-vaccine timepoint by prior commercial testing. Pre-versus post-vaccine endpoint titers compared using paired t tests. CD4 FS versus endpoint titers compared with Pearson’s correlation. A value of 1:49 was used for samples negative at the initial 1:50 dilution. ***p<0.001

## DISCUSSION

Respiratory viruses remain a major cause of morbidity and mortality in infants, and there is a critical need to interrogate adaptive immune responses in infants to improve the design of safe and highly effective vaccines. To that end, we examined the CD4 T-cell responses induced by primary SARS-CoV-2 mRNA vaccine series in infants, relative to those generated in adults. We found that infants had lower frequencies of Spike-specific CD4 T-cells that produced Th1 cytokines, particularly IFN-γ, relative to adults. This deficit translated to lower functionality / polyfunctionality as reflected in lower post-vaccination FS and PFS relative to adults. Further, the CD4 T-cell responses were higher in recipients who received mRNA-1273 vs. BNT162b recipients, though these differences were only significant in infants. In contrast to CD4 T-cell responses, Spike-specific post-vaccine IgG titers were similar between infants and adults.

Among infants, they were not modified by vaccine manufacturer or the presence of pre-vaccine maternal Spike-specific IgG. Together, these data emphasize that infants have lower CD4 T-cell induction following vaccination relative to adults, with the greatest impact on IFN-γ secreting subsets.

Infant T-cells are equipped to mount antigen-specific responses such as following vaccination, but their adaptive responses are altered compared to adults. Both T-cell function and memory response may be attenuated, which has been attributed to dendritic cell immaturity and restricted antigen internalization, processing, and presentation (29). Infant T-cells are highly proliferative but produce less IFN-γ and other Th1 cytokines than adult T-cells (30-32). For example, infants have previously been found to produce less IFN-γ than adults in response to measles or mumps vaccination (30, 31). Instead, immune stimuli may induce a predominant Th2 or Th17 response, although this difference has primarily been observed in infants younger than three months old (32-34). Although our observation of a deficit in CD4 T-cell production of IFN-γ amongst infants is consistent with the former of these observations, we did not find evidence of a shift toward a predominant Th2 or Th17 response, consistent with a recent study investigating T-cell responses to mRNA-1273 in children (19). In fact, the magnitude of Th2 response induced by vaccination was greater in adults than children. In contrast to infants, our adults had robust Th1 and to a lesser extent Th2 response to vaccination, consistent with previously described Spike-specific CD4 T-cell cytokine producing frequencies (20).

Both the mRNA-1273 and BNT162b mRNA vaccines are safe and effective for persons as young as 6 months of age. In this infant cohort, we found a stronger response to vaccination among infants who received mRNA-1273. Though the adult response was also slightly higher following mRNA-1273, this difference was not statistically significant. A potential explanation for the difference in CD4 T-cells responses we observed between mRNA-1273 and BNT162b is the difference in mRNA dose in the two vaccines despite the extra dose recommendation for BNT162b (infant vaccine: 25 µg/dose vs. 3 µg/dose; adult vaccines: 100 µg/dose vs 30 µg/dose, respectively (35, 36)). Within adults, mRNA-1273 has been associated with both higher Spike-specific endpoint titers and reduced risk of severe disease, particularly amongst the elderly (37-39). In contrast, no difference has been identified in Spike-specific IgG induced by the two vaccines in children aged 6 months to 5 years old (40). Further investigation into vaccine efficacy for infants and children is needed to discern if the accentuated CD4 T-cell response generated by mRNA-1273 leads to better disease control when infected.

Our study has several limitations. First, our cohort size was relatively small, but this was offset by the ability to collect sufficient volume of blood from infants to robustly characterize CD4 T-cell responses. Further, despite the size, the cohort had limited heterogeneity, and all of our participants were NC-IgG negative at both the pre- and post-vaccine time points, emphasizing that the effects we observed were secondary to vaccination alone and not hybrid immunity. Second, infant participants had their blood drawn a median of 4 weeks following the final vaccine dose, whereas adults were a median of 2 weeks post-vaccine at their final blood draw. Though the kinetics and durability of the T-cell response to SARS-CoV-2 vaccination remain understudied, particularly in pediatric populations, a previous study has identified stable COMPASS-derived spike-specific CD4 functionality scores for up to 12 months in unvaccinated children younger than 5 years old following infection (41). We additionally controlled for time of sampling post-vaccination in our multivariate models to account for this difference.

Together, our findings emphasize the importance of studying CD4 T-cell responses induced by vaccination, in addition to antibody titers, where CD4 T-cell responses may be particularly important in the control of infection and the prevention of severe disease. In addition, CD4 T-cell responses may be differentially impacted by age and vaccine manufacturer. Our data emphasize the need to specifically consider immune ontogeny in the design of novel pediatric vaccines. Future studies should determine whether the differences in CD4 T-cell responses we observed translate to differences in SARS-CoV-2 disease severity amongst vaccinated individuals.

## DATA AVAILABILITY

All primary data available in Supplementary Table 2

## FOOTNOTES

### Funding

Funding was provided by Burroughs Wellcome Fund (BWF CAMS 1017213; WEH), University of Washington, Department of Pediatrics (WEH), Seattle Children’s Research Institute (WEH), University of Washington Royalty Research fund (AK), the National Institute of Allergy and Infectious Diseases (K23 AI153390; AK), and the National Center for Advancing Translational Sciences of the National Institutes of Health (UL1 TR002319).

### Conflicts of Interest

MQP, QLY, JES, MG, EL, JKL, NKM, DNS, CS, and WEH have no conflicts of interest. JAE has received grant funding from AstraZeneca, GlaxoSmithKline, Pfizer, and Moderna and is a paid consultant for Abbvie, AstraZeneca, GlaxoSmithKline, Merck, Meissa Vaccines, Moderna, Pfizer, Shionogi, and Cidarra. AK is a co-investigator on a study funded by Pfizer, was an unpaid consultant for Pfizer and GlaxoSmithKline, and received a speaker’s honorarium from SCL Seqirus.

## ALT TEXT for FIGURES

Figure 1. Overview of vaccine schedule for infants and adults included in the study.

Figure 2. Summary of individual cytokines produced by CD4 T-cells following SARS2 mRNA vaccination in adults and infants with statistical comparison.

Figure 3. Polyfunctional analysis of cytokine production by CD4 T-cells following SARS2 mRNA vaccination in adults and infants using COMPASS algorithm. Includes analysis by vaccine manufacturer with statistical comparison.

Figure 4. Comparison of CD4 T cell responses to antibody responses following SARS2 mRNA vaccination in adults and infants with statistical comparison\.

**Supplementary Table 1.**
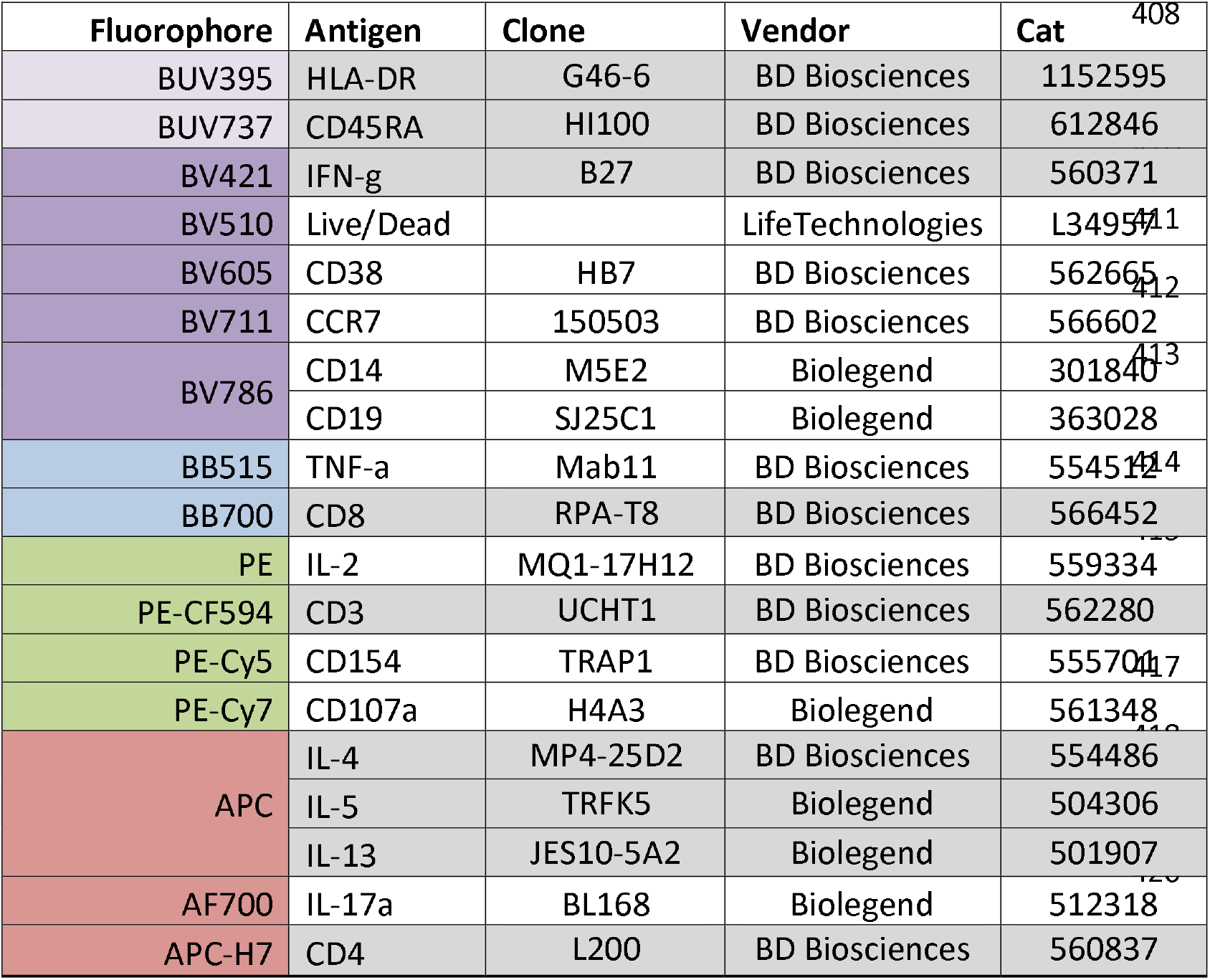
ICS flow cytometry panel.

## Notes

### Competing Interest Statement

Outside of this work, AK was an unpaid consultant for Pfizer and GlaxoSmithKline and received a speaker's honorarium from SCL Seqirus. AK is co-investigator on a study funded by Pfizer. JAE receives grant support to her institution from Merck, GlaxoSmithKline, AstraZeneca, and Pfizer and is a consultant for Abbvie, AstraZeneca, GlaxoSmithKline, Moderna, Pfizer, Sanofi Pasteur and Meissa Vaccines, Inc.

